# The safety and efficacy of Ultrasound Histotripsy and human pluripotent stem cell-derived hepatic spheroid implantation as a potential therapy for treatment of congenital metabolic liver disease: assessment in an immunocompetent rodent model

**DOI:** 10.1101/2024.11.08.622652

**Authors:** Hassan Rashidi, Amjad Khalil, Saied Froghi, Andrew Hall, Pierre Gelat, Brian Davidson, Alberto Quaglia, Nader Saffari

**Author notes:** These authors contributed equally (Hassan Rashidi, Amjad Khalil, Saied Froghi).

## Abstract

**Background:** Liver disease secondary to an inborn or genetic error of metabolism is a rare group of conditions often associated with chronic ill health and reduced survival. Curative treatment is mainly limited to liver transplantation with major long-term risks. Cell therapy is a promising alternative, but current approaches are ineffective.

**Aim:** To develop histotripsy, a non-invasive high intensity ultrasound procedure for liver tissue mechanical ablation, combined with hepatocyte stem cell implantation as a novel method of reversing liver failure from genetic disease. This study assessed the safety and feasibility of this approach in healthy rodents.

**Methods:** Under general anaesthesia, adult rats (n=12) underwent laparotomy and ultrasound histotripsy to the exposed liver. Around 1 million cells were injected into a single histotripsy cavity in each animal under direct vision (n=10) with two receiving only histotripsy without cell injection. On completion of cell implant haemostasis was secured, laparotomy incision closed, and the animals recovered. Groups of animals were terminated immediately and after 4 hrs, 8 hrs, 24hrs, 4 days and 7 days. Liver and vital organs were assessed for procedure related injuries and evidence of viable implanted cells by histology and immunohistochemistry.

**Results:** All animals successfully recovered, and no complication was observed throughout the study. Created cavities were successfully identified in histological analysis of rat. The presence of human cells was verified using anti-human nuclei antibody confirming successful implantation of liver organoids into decellularized cavities.

**Conclusions:** In this feasibility study, we demonstrated suitability of histotripsy to create decellularized cavities in liver parenchyma. In addition, feasibility of direct transplantation of undissociated liver organoids into the created cavities was demonstrated as a potential approach to treat inborn liver disease by creating nodules of healthy cells capable of performing loss metabolic function. Therapeutic efficacy of this approach will be evaluated in an upcoming study.

## Introduction

Congenital liver diseases are diverse group of genetic disorders that primarily affect liver function from birth. These conditions can lead to significant morbidity and mortality due to liver dysfunction resulted from accumulation of toxic metabolites (Scorza et al., 2014). Inborn metabolic disorders are rare but remain an important cause of liver disease (Vakili et al., 2024). Liver transplantation (LT) is one of the few effective treatments but is associated with a significant mortality in children (∼5% within the first year) and with a high risk of post-transplantation complications (up to 70%). In addition, there is a shortage of suitable donor organs for children, a risk of rejection and a requirement for life-long immunosuppression treatment which is associated with increased risk of sepsis, de novo cancer and renal failure (Khalil et al., 2023).

Alternative approaches have been sought to overcome limitations associated with LT including pharmacological interventions, gene therapy and cell therapy (Alwahsh et al., 2018, Ghasemzad et al., 2022). Cell therapy can be used as a curative option in metabolic disorders or as a bridge to transplantation (Khalil et al., 2023). Primary human hepatocytes (PHHs) have been considered as the optimum cell source for transplantation but represent a limited resource (Zeilinger et al., 2016).

Human pluripotent stem cells (hPSCs) possess unique features including the ability to be expanded indefinitely and the potential to be differentiated into any cell type found in the human body, making them an attractive source for cell therapies (Alhaque et al., 2018). To this end, various approaches have been developed to generate functional hepatocytes from hPSCs as a renewable source (Hay et al., 2007, Agarwal et al., 2008, Takebe et al., 2013, Hu et al., 2018, Rashidi et al., 2018, Tomaz et al., 2022).

Several approaches have been used for cell implantation into the liver or ectopically into other organs such as the spleen, peritoneal cavity and lymph nodes (Strom et al., 1997, Dhawan et al., 2010, Komori et al., 2012, Dwyer et al., 2021). Endovascular infusion, particularly into the portal vein, has been the preferred and most common route for hepatocyte transplantation in a clinical setting (Gramignoli et al., 2015). Embolism of terminal radicals of the portal system following transfusion of hepatocytes and subsequent vascular permeabilization has been suggested as the potential mechanism of cell integration. The embolism leads to portal hypertension, triggering the release of vasculature permeabilization cytokines from Kupffer and stellate cells that facilitate integration of transplanted cells into the liver parenchyma (Gupta et al., 1999). The engraftment efficiency is low at 5 to 30% as most of the non-integrated hepatocytes that remain wedged in the portal vessels are cleared by macrophages within 24 hrs (Joseph et al., 2002). The engraftment efficiency can be enhanced following preconditioning treatment of the liver by disruption of the sinusoidal endothelium, administration of vasodilators, inhibition of macrophage function, and irradiation though most of these strategies are not clinically acceptable (Malhi et al., 2002, Slehria et al., 2002, Joseph et al., 2002, Yamanouchi et al., 2009). Therefore, development of novel cell engraftment methodologies is a necessity in the field of liver cell therapy.

Ultrasound histotripsy can be used to mechanically fractionate soft tissues with a high degree of precision and has been used for non-invasive ablation of solid tumours (Aubry et al., 2013, Khokhlova and Hwang, 2016). To develop an efficient and targeted transplantation methodology, a protocol known as boiling histotripsy was previously used by our group to create well-defined decellularized cavities in rat liver without any noticeable thermal damage (Pahk et al., 2016).

The aim of this study was to use our previously developed US histotripsy protocol with the inclusion of a pulse-echo ultrasound transducer to generate a liver cavity in which to transplant undissociated hPSC-derived 3D Hepatospheres (3D-Heps) and to determine the feasibility and safety of this approach as a potential treatment for congenital metabolic liver disease.

## Materials and Methods

### Animals

Male Sprague–Dawley rats, 6–8 weeks old and weighing 430-530 g, were obtained from the Charles-River Laboratories UK Ltd (Margate, Kent, UK). The animals were housed in a temperature-controlled room (19-24°C), with a relative humidity of 40-70% and alternate light/dark conditions. They were given standard laboratory rodent chow. All animal experiments were conducted according to the Home Office guidelines under the UK Animals and Scientific Procedures Act 1986. All experiments were done under isoflurane general anaesthesia.

### Maintenance of human induced pluripotent stem cells (hiPSCs)

An inhouse generated hiPSC line (GOS101B) was used in this study. GOS101B was cultured on laminin 521 (BioLamina) coated plates in serum-free mTeSR^TM^1 medium (STEMCELL Technologies) as previously described (Rashidi et al., 2022). The cell lines were propagated in antibiotic free medium and were monitored regularly for mycoplasma infection using MycoAlert^TM^ PLUS Mycoplasma Detection Kit (Lonza).

### Generation of self-aggregated hiPSC spheroids

A previously published protocol was used to generate 3D Heps from hPSCs (Lucendo-Villarin et al., 2019). Initially, Agarose microplates were fabricated in 256-well format using the MicroTissues® 3D Petri Dish® mould (Sigma Aldrich) following the manufacturer’s instructions and transferred to 12-well plates (Corning). hPSCs scaled up on laminin-coated plasticware, were incubated with 1 ml of Gentle Dissociation Buffer (STEMCELL Technologies) for 7 min at 37°C. Following this, single cell suspensions were prepared by pipetting the buffer up and down gently. The cell suspension was centrifuged at 0.2 relative centrifugal force (rcf) for 5 min, discarded the supernatant and resuspended in mTeSR1^TM^ supplemented with 10 µM Y-27,632 (Cayman Chemical Company) at a density of 2.0 × 10^6^ live cells/ml. The agarose microplates were seeded by transferring 190 µl of cell suspension. After 2–3 h, 1 ml mTeSR1^TM^ supplemented with 10 µM Y-27,632 was gently added to each well of the 12-well plate and incubated overnight to allow formation of self-aggregated spheroids as described elsewhere (Lucendo-Villarin et al., 2019).

### Differentiation into 3D Heps

Differentiation was initiated a day after generation of hiPSC spheroids by replacing mTeSR1^TM^ with endoderm differentiation medium which consisted of RPMI 1640 containing 1 × B27 (Life Technologies), 100 ng/mL Activin A (PeproTech), and 50 ng/mL Wnt3a (R&D Systems). The medium was changed every 24 h for 72 h. On day 4, the endoderm differentiation medium was replaced with a hepatoblast differentiation medium. The medium consisted of knockout (KO)-DMEM (Life Technologies), knockout serum replacement (KOSR - Life Technologies), 0.5% Glutamax (Life Technologies), 1% non-essential amino acids (Life Technologies), 0.2% b-mercaptoethanol (Life Technologies), and 1% DMSO (Sigma), and was renewed every second day for a further 5 days. On day 9, the medium was replaced by a hepatocyte maturation medium HepatoZYME (Life Technologies) containing 1% Glutamax (Life Technologies), supplemented with 10 ng/ml hepatocyte growth factor (HGF; PeproTech) and 20 ng/ml oncostatin m (PeproTech). On day 21, cells were cultured in a maintenance medium consisting of William’s E media (Life Technologies), supplemented with 10 ng/ml EGF (R&D Systems), 10 ng/ml VEGF (R&D Systems), 10 ng/ml HGF (PeproTech), 10 ng/ml bFGF (PeproTech), 10% KOSR, 1% Glutamax, and 1% penicillin–streptomycin (Thermo Fisher Scientific).

### Immunofluorescence staining

Prior to implantation, 3D Heps were fixed in ice-cold methanol for 30 min, washed in PBS, and embedded in agarose before embedding in paraffin to obtain sections of 4 µm thickness. Antigen retrieval was performed by heating dewaxed and rehydrated sections in 1 × Trisodium Citrate buffer solution for 15 minutes in a microwave oven using defrosting option to avoid boiling of the buffer. Washed slides were used for subsequent staining to characterise generated cells.

To perform immunostaining, stem cell-derived tissue was blocked with 10% BSA in PBS-tween (PBST) and incubated with primary antibody overnight at 4 °C and detected using species-specific fluorescent-conjugated secondary antibody (Alexa Flour 488/Alexa Flour 568/Alexa Flour 680; Invitrogen). Sections were counterstained with DAPI (4′,6-diamidino-2-phenylindole) and mounted with ClearMount^TM^ mounting solution (Invitrogen).

An extended staining protocol was developed and optimised to stain whole mount 3D Heps. Briefly, MeOH fixed spheroids were washed and rehydrated in PBS. Following overnight blocking in 10% BSA in PBST, the spheroids were incubated with primary antibody at desired concentration (Supplementary Table 1) with gentle agitation at 4 °C overnight followed by 8 washes with 0.1% PBST (each wash 1 h) under gentle agitation at room temperature (RT). Secondary antibodies were incubated at 4 °C overnight and washed as described above. The nucleus was counterstained with DAPI (Life Technologies) before images were acquired using a Zeiss LSM 880 upright multiphoton confocal microscope.

### Cytochrome P450 Assay

3D Heps derived from hiPSCs were incubated for 1.5, 4 and 24 hours, with the luciferin conjugated specific CYP3A (1:40) substrate (P450 P-Glo Luminescence Kit, Promega) at 37°C. The Luciferin detection reagent was reconstituted by mixing the buffer into the bottle containing the lyophilised Luciferin detection reagent. For measurement, in a white 96 well plate 50 µl of the supernatant sample mixed with 50 µl of the detection reagent was added and incubated at room temperature in the dark for 20 minutes. The data was collected using a luminometer (GloMax Navigator, Promega). For data analysis, units of activity were measured as relative light units per ml per mg protein (RLU/ml/mg) as determined by the BCA assay.

### Gene expression analysis

Quantitative real-time polymerase chain reaction (qRT-PCR) was performed to demonstrate expression of the liver-specific genes HNF4A, ALB and CYP3A4. RNA was extracted from 3D Heps using the Direct-zol RNA Miniprep kit (Zymo Research). RNA quantity and quality were assessed using a Nanodrop system. Following this, cDNA was amplified using the RevertAid First Strand cDNA Synthesis kit (Thermo Fisher Scientific) following the manufacturer’s instruction. qPCR was performed with TaqMan Fast Advance Mastermix and primer pairs listed in Supplementary Table 2 and analysed using a CFX96^TM^ Real-Time PCR System (Bio-RAD). Gene expression was normalised to glyceraldehyde 3-phosphate dehydrogenase (GAPDH) and expressed as relative expression over 3D aggregate on day 0 of differentiation as control sample. qPCR was performed in triplicate and data analysis was performed using manufacturer software and graphs were prepared using GraphPad PRISM software (version 10.1.2).

### Enzyme-Linked Immunosorbent Assay (ELISA)

3D Heps derived from hiPSCs were incubated with maintenance medium for 24 hours at 37 ֯C in 5% (v/v) CO_2_, 95% (v/v) O_2_. The supernatants were collected after 24 hours. As markers of hepatocyte functionality albumin (ALB) and alpha-fetoprotein (AFP) levels were measured, using human ALB and AFP DuoSet ELISA kits (R&D System).

To detect human AFP and ALB in transplanted animals, 100 µl of plasma samples were loaded into pre-coated ELISA plate in duplicate followed by 1-hour incubation at room temperature as per manufacturer’s instructions. Following this step, microwells were washed with working wash solution for four times and diluted anti-human AFP or ALB HRP conjugated was added to the microwells. The wells were incubated at room temperature for 30 minutes. Microwells were washed with working wash solution five times, and the substrate for the HRP enzyme TMB, was added and incubated for 15 minutes in the dark. Following this step, the stop solution was added to each well and the plates luminous activity was measured at 450 nm with a reference wavelength of 630 nm using a FLUOstart Omega plate reader (BMG LabTech, Germany). Plain tissue culture media which was incubated for 24 hours at 37°C was employed as the negative control. For data analysis, the collected data was normalised per ml per mg protein as measured by the BCA Assay (Pierce, UK).

### Development of ultrasound histotripsy treatment Protocol

A 2 MHz single element spherical section transducer (Sonic Concepts H-148, Bothell, WA, USA) with an aperture size of 64 mm, a radius of curvature of 63.2 mm and a 22.6 mm central aperture was used with a transparent coupling cone (Sonic Concepts, C-101, Bothell, WA, USA) filled with degassed, de-ionised water. The cone was coupled directly to the animal’s exteriorized liver via a 12 μm-thick acoustically transparent polyethylene (Mylar) film (PMX 980, HiFi Industrial Film, Stevenage, UK). The transducer was driven by two function generators (Agilent 33220A, CA, USA) connected in series to a linear radiofrequency (RF) power amplifier (ENI 1040L, Rochester, NY, USA). The first function generator was set to generate pulses of a 1 Hz rectangular wave with 1% duty cycle. This triggered the second function generator, which outputted a 2 MHz sinusoidal wave into the RF power amplifier. Therefore, the drive signal into the amplifier was 30 to 35 pulses of 10 ms duration containing 20,000 cycles each. A power meter (Sonic Concepts 22A, Bothell, WA, USA) was connected between the RF amplifier and the ultrasound transducer, and the electrical power supplied to the transducer was monitored to be approximately 150 W. The pulse-averaged electrical powers was 1.5 W (calculated using P_averaged_ = P_peak_ × duty cycle). Assuming a nominal electrical to the acoustic power conversion efficiency of 85% (Sonic Concepts, Bothell, WA, USA) the acoustic peak positive (P+) and negative (P-) pressures at the ultrasound beam focus in liver tissue were P+ = 76.7 MPa and P- = –13.4 MPa, obtained by numerically solving the Khokhlov-Zabolotskaya-Kuznetsov (KZK) parabolic nonlinear wave propagation equation for our input parameters using the HITU Simulator v2.0 (Soneson, 2017). These pressure distributions are shown in Figure 1.

**Figure 1:**
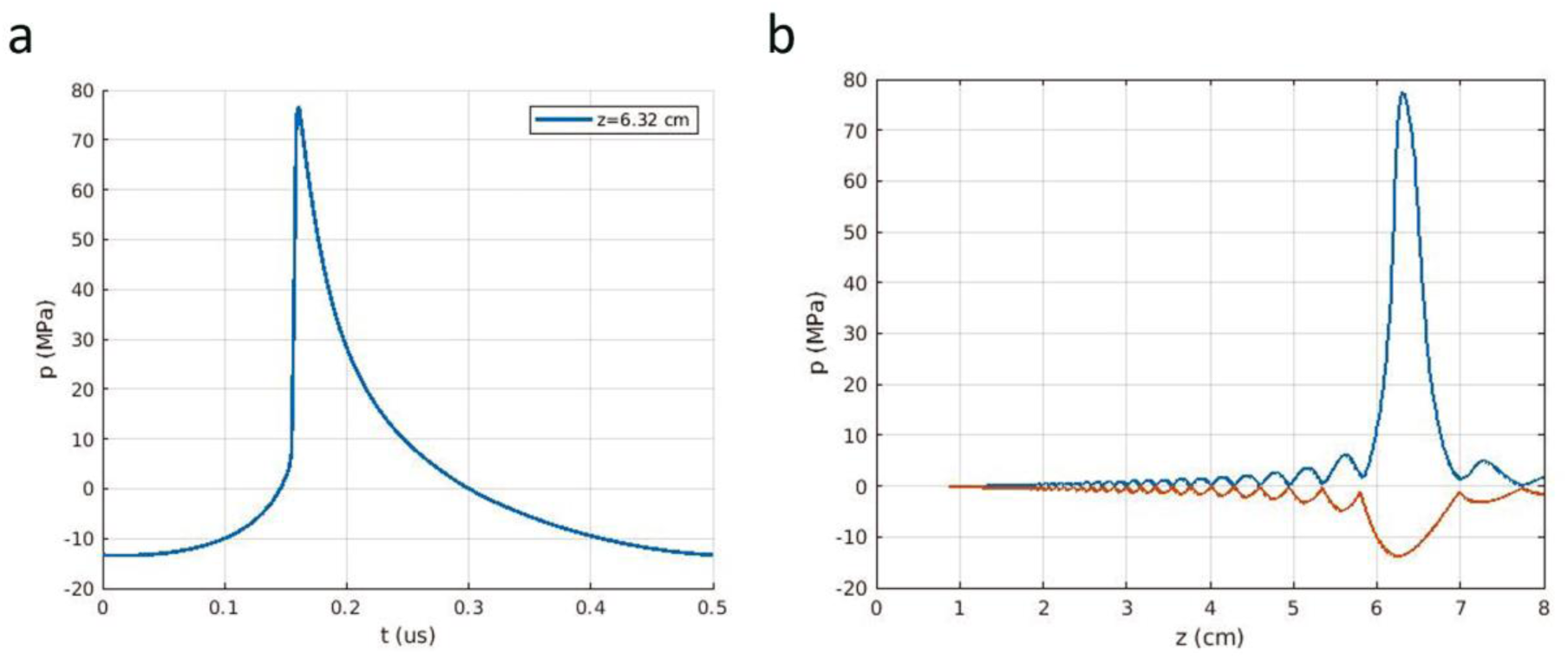
Numerically simulated ultrasound waveforms (a) at the geometrical focus of the transducer in liver tissue and (b) the axial distribution of the positive and negative peak pressure values.

To ensure correct and consistent coupling between the cone and the liver surface, we inserted a pulse-echo transducer (Olympus V312, 10 MHz centre frequency, 6 mm element diameter, unfocused) in the central open aperture of the Sonic Concepts H-148 transducer, as shown in figure 2. This transducer was connected to a Panametrics 5055PR pulser-receiver for the excitation and detection of the returned echoes from the surface of the liver.

**Figure 2:**
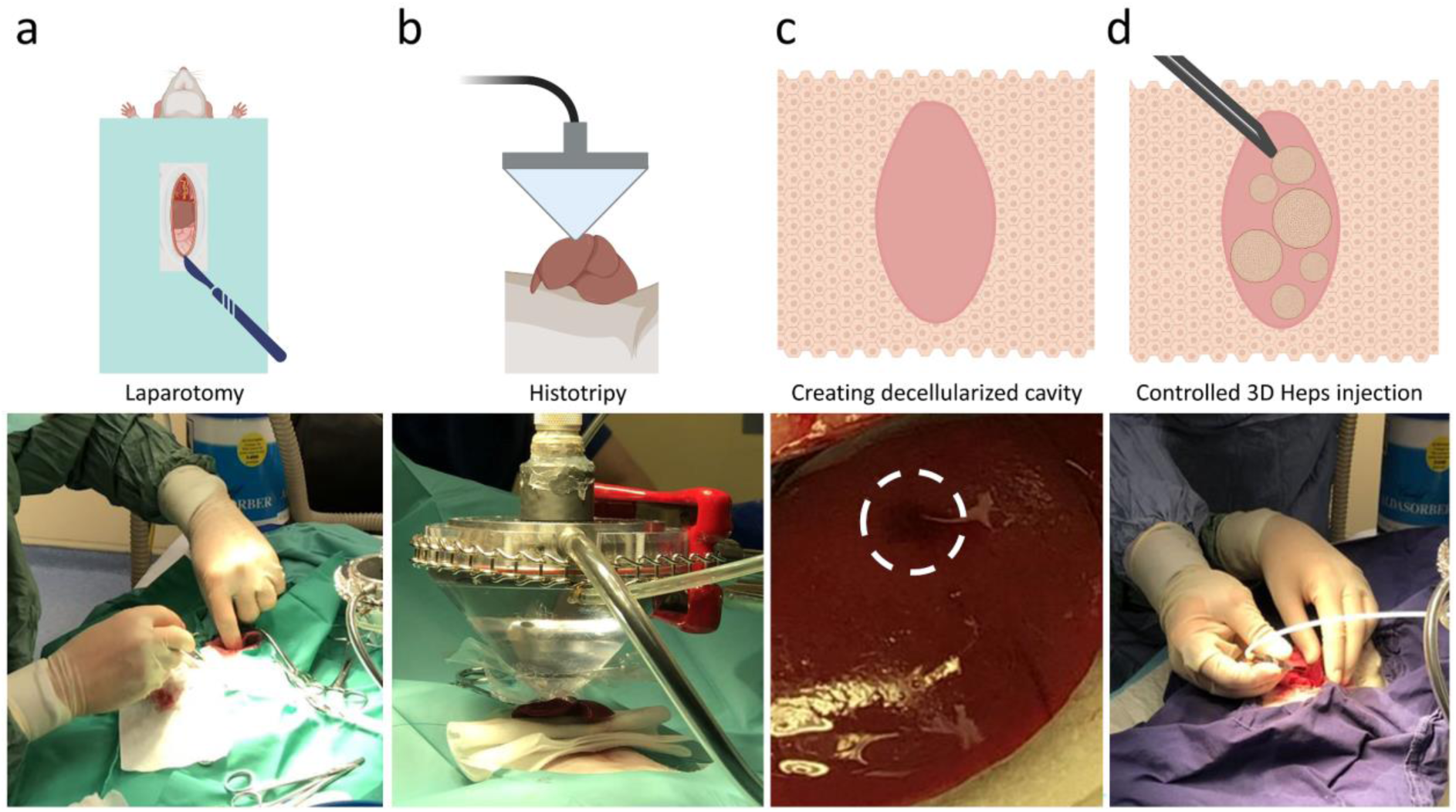
Histotripy decellularization and cell transplantation. **a) Laparotomy to allow exteriorisation of the liver.** **b) The histotripsy transducer was adjusted to touch the liver capsule lightly before initiating the protocol.** **c) Creation of the cavity was confirmed visually following observation of haematoma under liver capsule (indicated with white dashed circle).** **d) 3D Heps were administered using a syringe pump to permit controlled injection of the designated volume to avoid rupturing the liver.** **(Illustration made using Biorender.com)**

### Surgical procedure

Male Sprague Dawley Rats (n=12) were individually placed in an anaesthetic induction chamber. Flow of Isoflurane with oxygen was gradually increased until anaesthesia was induced. The animal was removed from the chamber, with anaesthesia maintained via a close-fitting mask. Animals were placed on a warm pad in theatre, abdominal hair clipped, and skin cleaned with chlorhexidine. Two drops of 2% lidocaine were injected subcutaneously followed by an upper midline laparotomy incision. Skin and fascia were retracted using 4/0 prolene sutures and weighted down with mosquito forceps. The right median lobe of the liver was mobilised preferentially and placed on sterile gauze, outside the abdominal cavity producing an air gap to produce a larger ultrasound **reflection** for measuring the thickness of the liver lobe.

The histotripsy probe was then lowered onto the exposed liver, so the exit aperture of the coupling cone was in contact with the exposed area of the liver. The histotripsy parameters are mentioned in supplementary table 1. Once the histotripsy procedure was complete, the experimental group received 200ul of cell suspension in saline buffer injected into the created cavity whilst the control group received histotripsy only (see figure2). To perform injection under control manner, a syringe pump was used to inject the volume at 1.2ml/min. A haemostatic gelatin matrix, containing thrombin (SURGIFLO®) was used to cover the cavity site in 3D Hep-implanted animals to prevent bleeding and leakage of implanted cells and it was applied on non-injected controls to keep procedure consistent between control and test groups. All animals underwent a single histotripsy procedure except animal 11 which underwent a second procedure as the presence of a cavitation haematoma was not observed after the initial sonification on the right median lobe. Following a check of haemostasis the abdomen was closed in layers with interrupted vicryl 3/0 to the fascia and prolene 3/0 to the skin. Veterinary Wound Powder and Opsite were administered to the closed wound sites.

Non-recovery animals were terminated immediately post procedure with cardiac puncture for venous sampling followed by intra-cardiac overdose with Sodium Pentobarbitone. Immediately following termination livers were harvested and immersed in formaldehyde solution, prior to paraffin fixation. Recovered animals were taken back to the animal accommodation area after a period of observation and received regular analgesia. The figure below outlines key procedures (Figure 2).

To examine creation of the cavity, cell transplantation and early onset of tissue repair, we sacrificed the animals immediately post-histotripsy (n=2; only histotripsy control), and immediately (n=2), 4 hrs (n=2), 18 hrs (n=2), 24 hrs (n=2), 4 days (n=1) and 7 days (n=1) post-3D Heps transplantation

### Histology

Each liver was dissected into 3-5 slices and fixed for at least 24 h in 10% neutral buffered formalin solution (pH7.4) at room temperature (RT). Samples were subsequently processed into Formalin Fixed Paraffin Embedded (FFPE) blocks for preservation.

For each block, we took a set of 7 serial sections of 4um thickness starting with the most superficial aspect and stained the first and last of the set of serial sections with Haematoxylin and Eosin (H&E). Sections were dewaxed in xylene and hydrated graded alcohols before staining for 10 mins in Harris’s haematoxylin (VWR cat no. 10047107). The slides were rinsed in tap water before differentiation in 1% acid (HCl) 70% alcohol (aqueous) before blueing in tap water for 5 mins. The slides were counterstained with Eosin (VWR cat no.10047101) for 5mins then washed in running tap water for 1 minute. The slides were then rapidly dehydrated, cleared and then mounted in DPX (VWR cat no 06522-500ML).

The H&E-stained slides were reviewed, if no cavity was seen the block was trimmed 100um deeper and another 7 serial sections were taken with the first and last being stained for H&E again for review. This procedure was repeated up to 14 times in order to identify any cavities throughout the tissue. Brightfield images were taken using a Hamamatsu Nanozoomer slide scanner.

### Immunohistological analysis

When a suspected cavity was found the reserved unstained serial sections between the stained H&E sections were stained with IHC Anti-Human Nucleoli antibody (NM95) - Nucleolar Marker for the detection of human nuclear antigen. Sections were dewaxed in xylene and hydrated in graded alcohols. Antigen retrieval was carried out (20 mins microwave at 640W in 1L of pH6.0 citrate buffer) before endogenous peroxidases were blocked (Bloxall - Vector Laboratories). Sections were incubated in 2.5% Horse serum (ImmPRESS Enzyme Polymer - Vector Laboratories) for 5 minutes and then incubated for 1 hour at room temperature (RT) in NM95 primary antibody (ab190710 16 - Abcam) at 1:1500 in TBS. Primary antibody binding was detected by incubating for 25 minutes in polymer solution (ImmPRESS Enzyme Polymer - Vector Laboratories) and developed with 3,3′ di-amino-benzidine - DAB (ab64238 - Abcam). Slides were counter-stained in Mayer’s Haematoxylin before they were dehydrated, cleared and mounted. Brightfield images were taken using a Hamamatsu Nanozoomer slide scanner.

### Statistical analysis

Data were analysed by GraphPad Prism (version 9.0). Student’s t test or Mann– Whitney tests were used to determine significant differences, which was set at P < 0.05. We did not use statistical methods to predetermine sample size, there was no randomization designed in the experiments, and the studies were not blinded.

## Results

### Generation of hPSC-derived 3D Heps and functional characterisation

Aggregated hPSCs were differentiated toward the hepatocyte lineage using a stepwise differentiation protocol described previously (Rashidi et al., 2018; Figure 3a). To produce homogenous three-dimensional spheres (3D Heps) of defined size, the agarose multi-well plate technology was employed to form hPSC aggregates. By day 30 of differentiation, 3D Heps exhibited typical hepatic morphology with distinctive cell borders which was maintained until harvesting the organoids for transplantation on day 60 of differentiation (Figure 3a). Differentiation to hepatic lineage was confirmed following detection of liver-specific genes by qPCR (Figure 3b). We also measured the level of AFP and ALB secretion using ELISA (Figure 3c) and CYP3A4 metabolic activity (Figure 3d). Finally, we performed immunostaining analysis of hepatic markers such as hepatocyte nuclear factor 4 alpha (HNF4A), AFP, ALB and CYP3A4 (Figure 3e).

**Figure 3:**
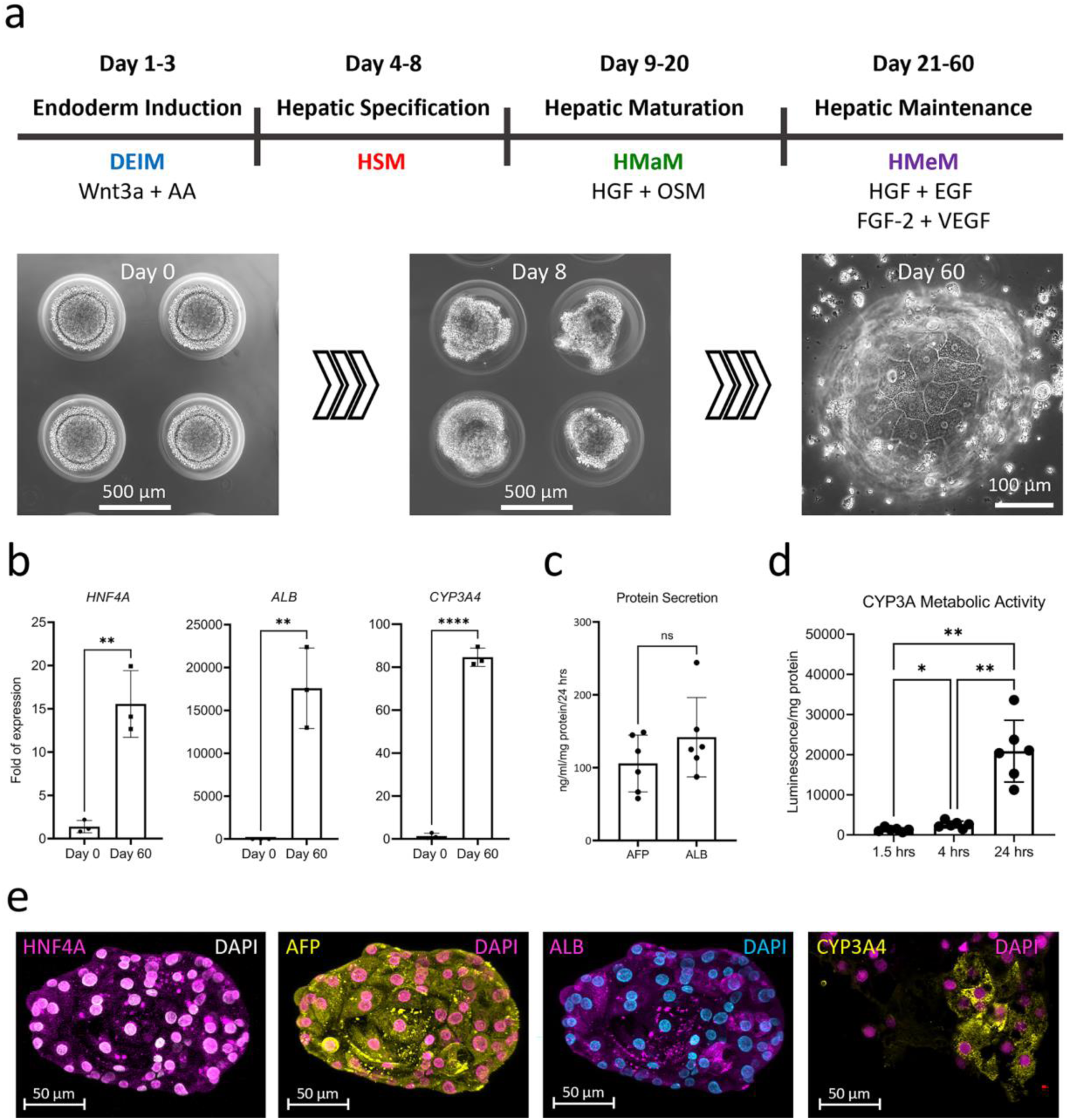
3D Heps characterisation. **a) A stepwise differentiation protocol was used to generate 3D Heps from hiPSCs.** **b) Expression of liver-specific genes HNF4A, ALB and CYP3A4.** **c) ELISA was used to measure AFP and ALB secretion by generated 3D Heps.** **d) Bioluminescence assay was used to assess CYP3A4 metabolic activity of 3D Heps.** **e) Immunostaining was performed to detect liver-specific proteins.** **(Abbreviations: DEIM, Definitive Endoderm Induction Medium; HSM, Hepatic Specification Medium; HMaM, Hepatic Maturation Medium; HMeM. Hepatic Maintenance Medium)**

### Termination of recovered animals

Recovered animals were terminated immediately after histotripsy and cell implantation (n= 2) and at 4 (n=2), 18 (n=2), 24 (n=2), 96 (n=1) and 168 hrs (n=1) post cell-implantation. Animals were terminated by placing the animals in an anaesthetic induction chamber, followed by a gradual increase in the flow of Isoflurane. Once anaesthetised a cardiac puncture was performed for venous sampling followed by intra-cardiac overdose with Sodium Pentobarbitone. Once death was confirmed a postmortem laparotomy and sternotomy were performed with lateral subcostal extension on each animal. Livers were preferentially harvested and immersed in formaldehyde solution, prior to paraffin fixation.

### Findings at postmortem

On macroscopic examination, animals terminated within 24 hours (Test 3 - 8) were found to have clean and healthy incision wounds, a single cavity within the liver (marked with SURGIFLO®) with minimal adherence to the diaphragm and abdominal wall. The livers appeared healthy, warm and well perfused. The remaining peritoneum and abdominal cavity appeared unremarkable, with no macroscopic, adhesions, haematoma, scarring, inflammation, injury or compensatory hypertrophy.

Animals terminated after 24 hours (test 9 & 10) had superficial wound dehiscence in the epigastric region of their laparotomy incisions, with the liver densely adhered to the incised abdominal wall and diaphragm with fibrous bands. The livers appeared healthy, warm and well perfused. The remaining peritoneum and abdominal cavity appeared unremarkable, with no macroscopic adhesions, haematoma, scarring, inflammation, injury or compensatory hypertrophy.

### Histological analysis

Created histotripsy sites were identified in one out of the two histotripsy control animals and four out of ten animals injected with 3D Heps (Figure 4). In the control animal, the histotripsy site consisted of a flask-shape decellularized area, filled with cell debris and red blood cells (Figure 4a-c) reaching a parenchymal depth of 4 mm below the liver surface and measuring 2 mm in maximum width. The histotripsy site reached the liver surface and was in contact with the Surgiflo gel plug without a visible interposed intact liver capsule at a point measuring 0.8 mm in diameter. The presence of 3D Heps in the form of cell aggregates was observed only in the livers implanted with cells (Figure 4d-p), of test animals 1, 2, 4, 8 and 10. The histotripsy sites showed a combination of necrosis and haemorrhage in animals 1, 2, 4 and 8, similar in appearance to the histotripsy site in the control animal. Three of these sites (animals 2, 4 and 8) showed an elliptical shape and were of variable length (2 mm, 1.2 mm and 3.6 mm) and width (0.5 mm, 0.5 mm and 1.4 mm, respectively), in the sections examined. The histotripsy site in animal 2 was located 2 mm from the liver capsule, which appeared intact (Figure 4d-f). A 1.2 x 0.5 mm separate subcapsular patch of necrosis and haemorrhage was also present. The histotripsy site in animal 4 was in contact with a 0.5 mm indentation of the liver capsule (Figure 4g-j). The histotripsy site in animal 8 was in contact with the liver capsule (see below). The histotripsy site in animal 1 was incompletely represented in the section examined, which also suffered from tissue tearing during processing and could not be measured or assessed in terms of relationships with the liver capsule (Supplementary Figure 2). Surgiflo gel plugs were identified in the samples from animals 8 and 10. The number of 3D Heps aggregates on prepared sections was variable. One aggregate was identified in animal 10, two in animals 2 and 4, and three in animals 1 and 8 (Figure 4d-p). The 3D Heps aggregate in animal 10 appeared to be wedged between the Surgiflo gel plug and the underlying intact liver parenchyma. Two of the three 3D Hep aggregates in animal 8 appeared to be wedged between necrotic parenchyma and the Surgiflo gel plug without a visible interposed intact liver capsule. The remaining eight 3D aggregates in animals 1, 2, 4 and 8 were positioned inside histotripsy sites. The 3D Heps appeared to be mostly viable histologically in test animal 1, 2, 4, and 8, but showed degenerative changes in animal 10. Focal basophilic material present in the 3D Heps in animal 1, 2, 4, and 8 represented probably fragmented nuclei. No inflammatory reaction was identified around the 3D Heps in animals 1, 2, 4. The histotripsy/injection sites in animals 8 and 10 contained infiltrates of leukocytes. In animal 10, the infiltrate was present between the Surgiflo gel plug, underlying liver parenchyma and a 3D Heps cluster and included a multinucleated giant cell (Figure 4n-p).

**Figure 4:**
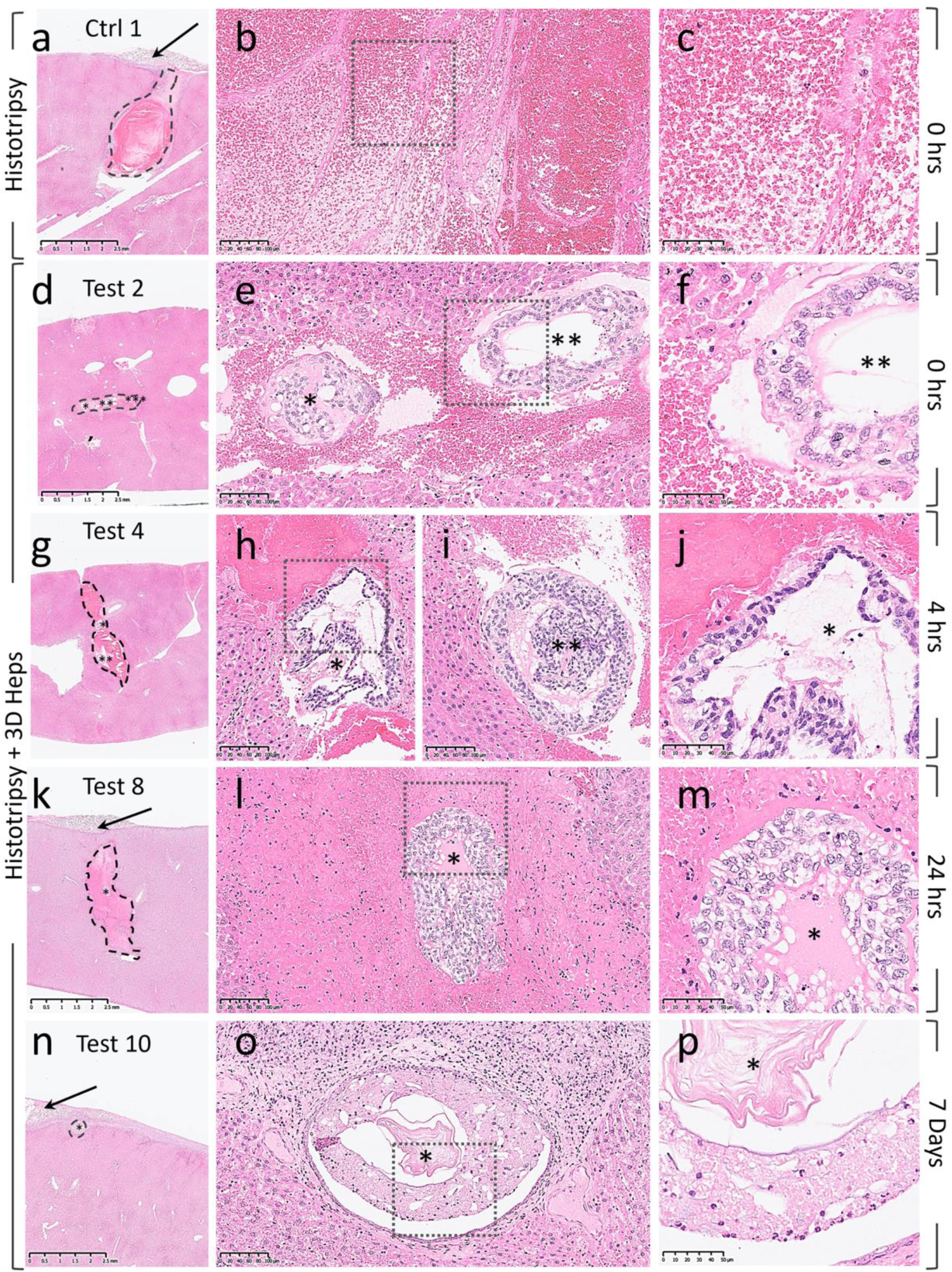
**Histological analysis. H&E staining was performed on sections obtained from rat livers following histotripsy (a-c) and 3D Heps transplantation (d-p). Implanted 3D Heps in Test Rat 10 (168 hrs post-implantation; n-p) was infiltrated with leukocytes (h-i). Cavities are marked with hatched black lines, 3D Heps with asterisks and SURGIFLO® with arrows.**

To confirm human origin, immunohistochemical analysis was performed using anti-human nuclei antibody. Stain-positive nuclei was observed only in samples injected with 3D Heps and where cell aggregates were presented (Figure 5).

**Figure 5:**
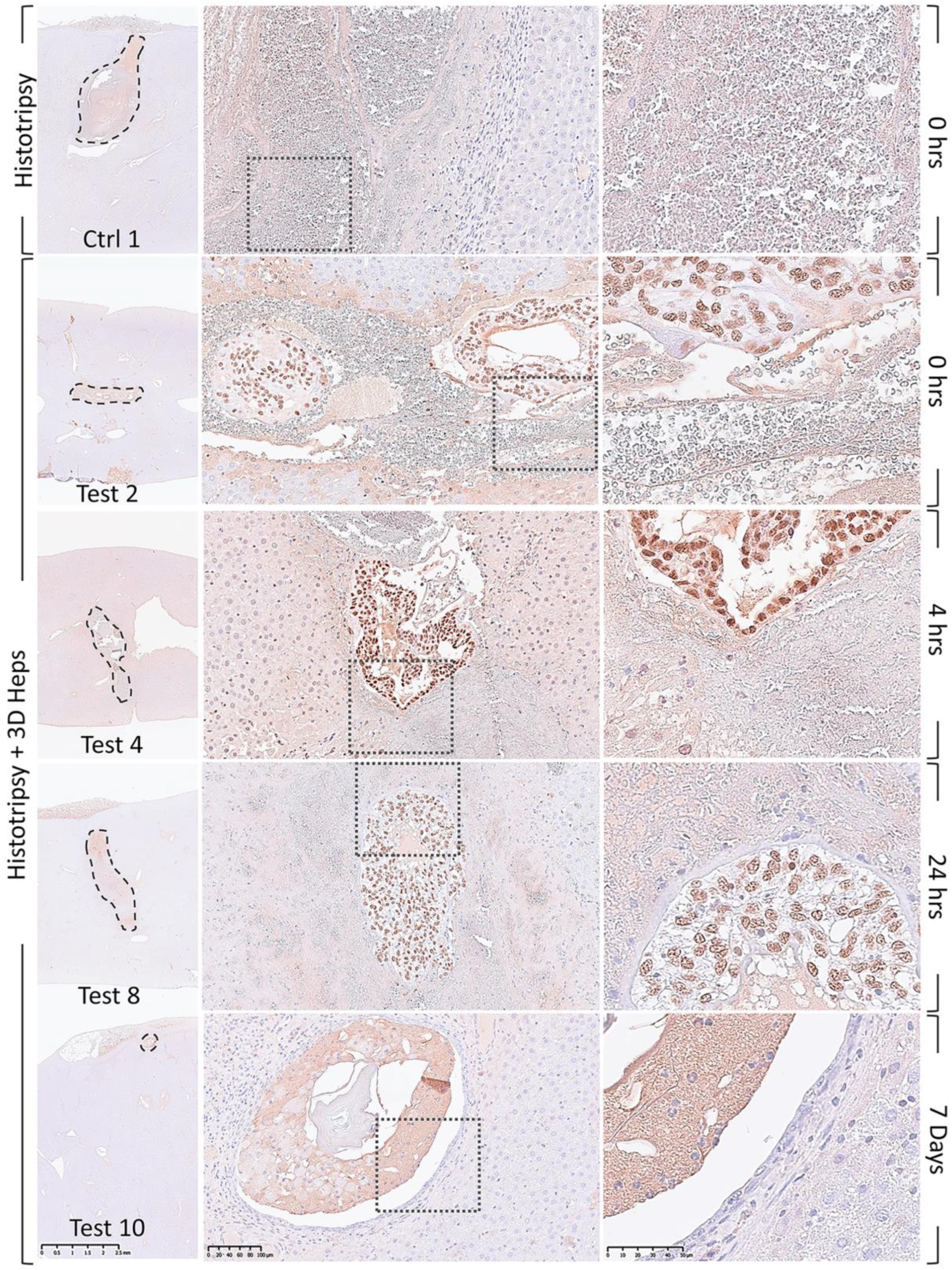
**The presence of human 3D Heps was validated using human anti-nuclei antibody. The immunostaining was negative for human nuclei in the aggregate identified in animal 10 possibly due to degenerative changes.**

### Functionality

We performed blood biochemistry on plasma collected from two controls and ten cell transplanted rats at the time of termination to detect human AFP and ALB. The collected plasma showed haemolysis in 8 samples of which 6 were heavily haemolysed. In addition, collected plasma was not sufficient in control rat 1 and cell transplanted rat 7 to run ALB ELISA (Supplementary Table 1). Human AFP was detected in the plasma of five out of the ten cell transplanted rats ranging from 0.016 to 1.883 ng/ml. ALB was detected in the plasma of 6 out of 9 transplanted rats ranging from 0.095 to 0.429 ng/ml (Figure 6; Supplementary Table 1). The detection and detected levels of AFP and ALB bore no correlation with collection time post-implantation. No human AFP or ALB was found in non-transplant control.

**Figure 6:**
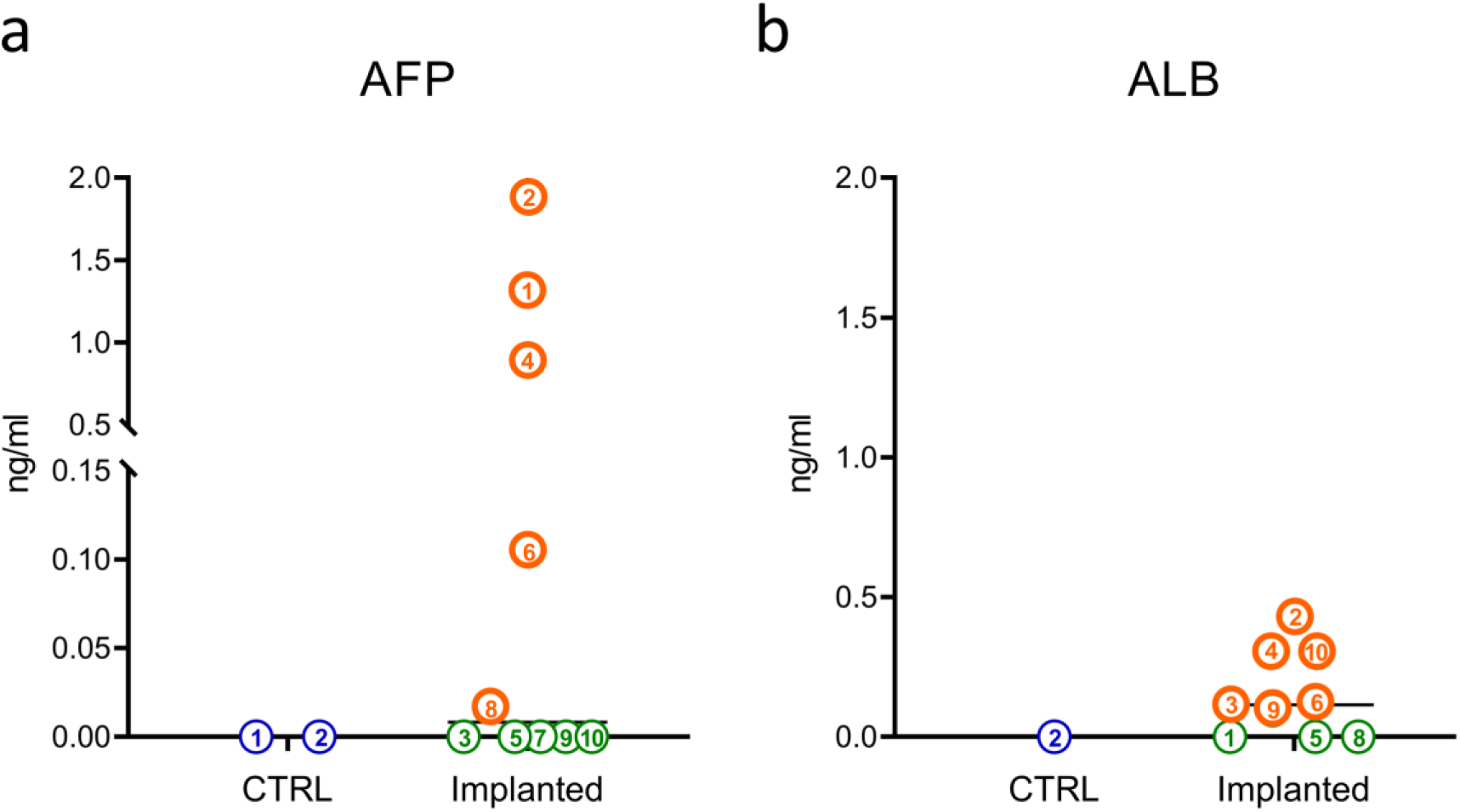
**Blood biochemistry analysis. Circulating human AFP was detected in 5 out of 10 cell transplanted rats. Human ALB was detected in 6 out of 9 cell transplanted animals. Plasmas positive for AFP and ALB are labelled in orange. The number inside circle shows designated number to rats in control and test groups. Collected serum from Ctrl Rat 1 and Test Rat 7 were not enough to include in ALB ELISA assay. Implanted Test Rats 1 & 2 sacrificed immediately post-implantation; 3 & 4, 4 hours post-implantation; 5 & 6, 18 hours post-implantation; 7 & 8, 24 hours post-implantation; 9, 96 hours and 10, 168 hours post-implantation.**

## Discussion

Cell therapy is a promising approach to treat liver disease, however, the lack of an efficient methodology for cell delivery remains a major bottleneck to the clinical translation of this modality. In this proof-of-concept study, we demonstrate the feasibility of producing a histotripsy cavity within the healthy liver and delivering hPSC-derived liver organoids (3D Heps) into the cavity as a novel approach to the treatment of liver failure in congenital metabolic liver disease.

To assess feasibility, we used rat as animal model. To allow access and establish direct contact with the ultrasound coupling cone, the liver was exteriorised outside the abdominal cavity. This approach facilitated liver stabilization as the rodents were not ventilated, with the gauze air trap beneath the liver aiding in reflecting the waves back to the pulse-echo transducer. However, this method posed challenges, including liver stretching that resulted in segment thinning, thereby elevating the risk of liver capsule puncture. Externalization of the liver also led to changes in perfusion, potentially causing venous congestion and subsequent elevation in capillary pressure, which may have contributed to the observed 3D Heps leakage in test rats 6 and 7 prior to the application of SURGIFLO®. The use of a bevelled needle may have contributed to leakage in these instances, either by penetrating too deeply, leading to bleeding and potential capsule damage, or by being too shallow, resulting in the leakage of 3D Heps from the syringe. In human, exteriorisation of the liver will not be required as the delivery of cells can be performed using ultrasound-guided percutaneous injection.

We used histotripsy to create cavities in the liver parenchyma and beneath the liver capsule, providing a suitable niche for cell transplantation. It has been shown previously that the dimensions of the cavity created depends on the ultrasound protocol selected (Pahk et al., 2019). In a previous publication, we reported the relationship between the dimensions of the cavity and the ultrasound parameters such as the output power and number of pulses employed, for a transducer of similar type (Pahk et al., 2017). The results presented as Figure 8 in Pahk et al. 2017, were obtained in an optically transparent liver phantom. We further confirmed the production of a cavity of the correct size for a given exposure setting in ex vivo chicken liver tissue before applying the same exposure conditions for our in vivo experiments. So as not to puncture the liver capsule, the axial length of the cavity is required to be smaller than 6 mm, which is the average liver lobe thickness in our male Sprague Dawley rats.

By including a pulse-echo transducer, we successfully evaluated liver lobe thicknesses in all animals except two, enabling precise selection of a sonication location to create a cavity within liver capsule boundaries and avoid puncturing the liver. Difficulty in assessing thickness of right median lobe in control rat 1 resulted in puncturing the capsule. Subsequently, we reduced the pulse count from 35 to 30 and sonicated rat 2’s left median lobe (the only animal to undergo left lobe sonication). Future studies will include real-time imaging for improved procedural monitoring.

Following histotripsy-mediated cell implantation, the animals were recovered in dedicated cages and exhibited expected normal behaviour, physical activity, water consumption, food intake, urination, and defecation patterns indicating the method safety. Apart from wound complications in test rats 9 and 10 that were kept for 96- and 168-hours post-implantation, the animals maintained a generally healthy appearance. Serum tests for overall rodent and liver health were not conducted during the recovery period. Potential reasons for wound complication include trauma (animals picking at it), poor surgical technique and infection.

Postmortem findings were as would be expected for the animals terminated within 24 hours. There was minimal adherence of cavities to adjacent diaphragm/ abdominal wall. SURGIFLO® a haemostatic agent was used on the cavities and has tissue adherence properties. Animals 9 & 10 had a firmer tissue reaction between likely sites of cavity formation and the abdominal wall / diaphragm. This could be expected post-histotripsy or as a result of xenograft response to injected 3DHeps. Histological analysis indicated that the resulting cavities were around 0.4 ± 0.1 mm beneath the liver capsule and were around 2.7 ± 0.7 mm in length and 1.0 ± 0.3 mm in width.

In a previous study from our group (Pahk et al., 2016), Matrigel^TM^ was used to improved retention of transplanted dissociated rat hepatocytes at the site of transplantation. Matrigel^TM^ is a solubilised basement membrane matrix secreted by Engelberth-Holm-Swarm mouse sarcoma cells which is rich in laminin and collagen-IV. As Matrigel^TM^ turns into gel at temperatures between 22 and 37 ֯C, it is a suitable protein carrier to improve cell retention at the site of transplantation. However, as an undefined animal product, its application in clinical practice risks the transfer of animal pathogens. In addition, mature hepatocytes are prone to cell death and epithelia-to-mesenchymal (EMT) transition and loss of functionality following dissociation (Cicchini et al., 2015). In this study, we transplanted undissociated hiPSC-derived 3D liver organoids (3D Heps) to circumvent EMT and cell death. However, the size of 3D Heps at around 200-300 μm in diameter made it more difficult to implant in comparison to dissociated hepatocytes. Creating small cavities of these sizes beneath liver capsule had imposed difficulties in identification of the generated cavities and implantation of undissociated 3D-Heps with accuracy. We successfully identified 5 out of 10 generated cavities and confirmed presence of implanted human 3D Heps using anti-human nuclei antibody. The presence of functional human cells was supported by the finding of human AFP and ALB only in the cell transplant animals. The detection of human AFP and ALB cannot be considered as a sign of viability and integration of implanted cells as both proteins are relatively stable and can be released from damaged cells post-implantation. Successful integration and long-term functionality should be studied in an immunocompromised animal model to prevent rejection of transplanted cells post-implantation. Our set of experiments is not sufficient to characterise the host response to the creation of cavities and cell implantation, which would be the subject of further studies.

In summary, we have obtained encouraging early feasibility data on histotripsy-mediated cell transplantation as a potential method of cell therapy for some of the congenital metabolic liver disease. While demonstrating short-term safety of the methodology, ongoing work will focus on long-term cell integration and functionality and assessment of the therapeutic efficacy. Low thickness of rat liver imposes limitation in number and size of the cavities that can be created and as a result the number of cells that can be transplanted; however, this would not be an issue following translation of the technique into human as larger cavities can be generated to transplant larger number of cells. In addition, the procedure can be performed percutaneously in human using ultrasound-guided cell implantation to avoid laparotomy and making the procedure less invasive.

## Acknowledgements

This work is supported by 2020\100261 Seedcorn Award from Rosetrees Trust. In addition, Hassan Rashidi is funded by the NIHR GOSH BRC. The views expressed are those of the author(s) and not necessarily those of the NHS, the NIHR or the Department of Health.

---

**Supplementary Figure 1:**
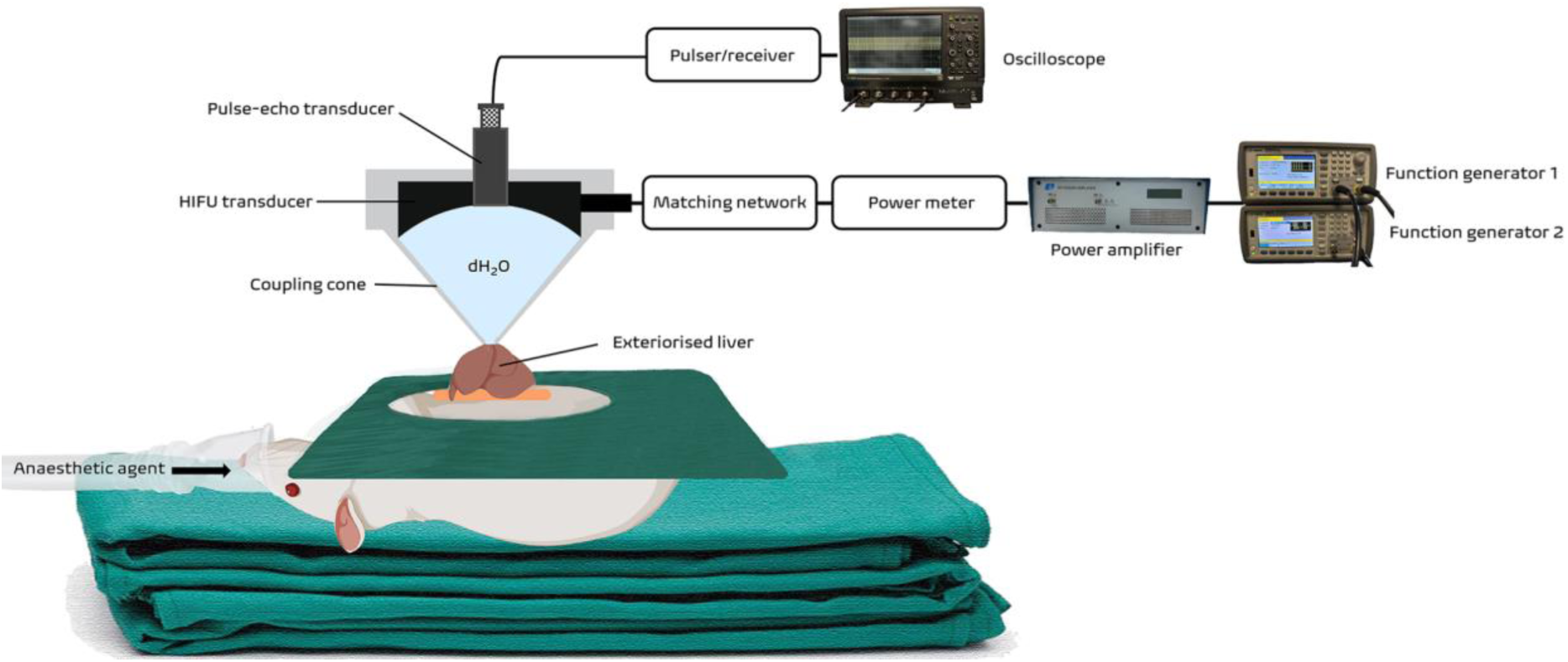
**Schematic of histotripsy-mediated cell transplantation. Due to anatomical location and size of the liver in the rat, it was exteriorised following laparotomy before performing histotripsy to create decellularized cavity.**

**Supplementary Figure 2:**
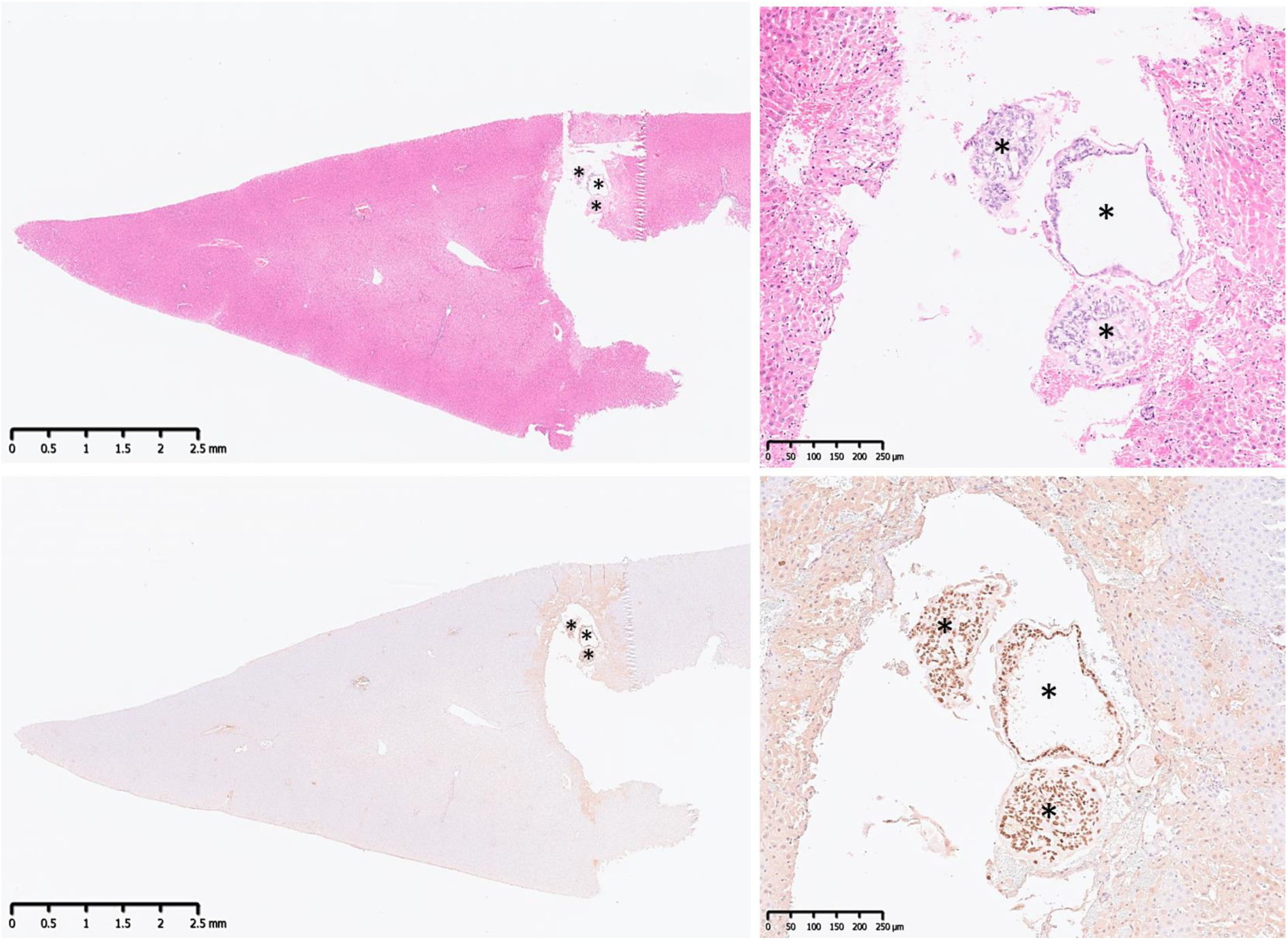
**Histological and immunohistochemistry analysis of sections obtained from Test 1 showing 3 implanted 3D Heps inside the cavity.**

**Supplementary Table 1:**
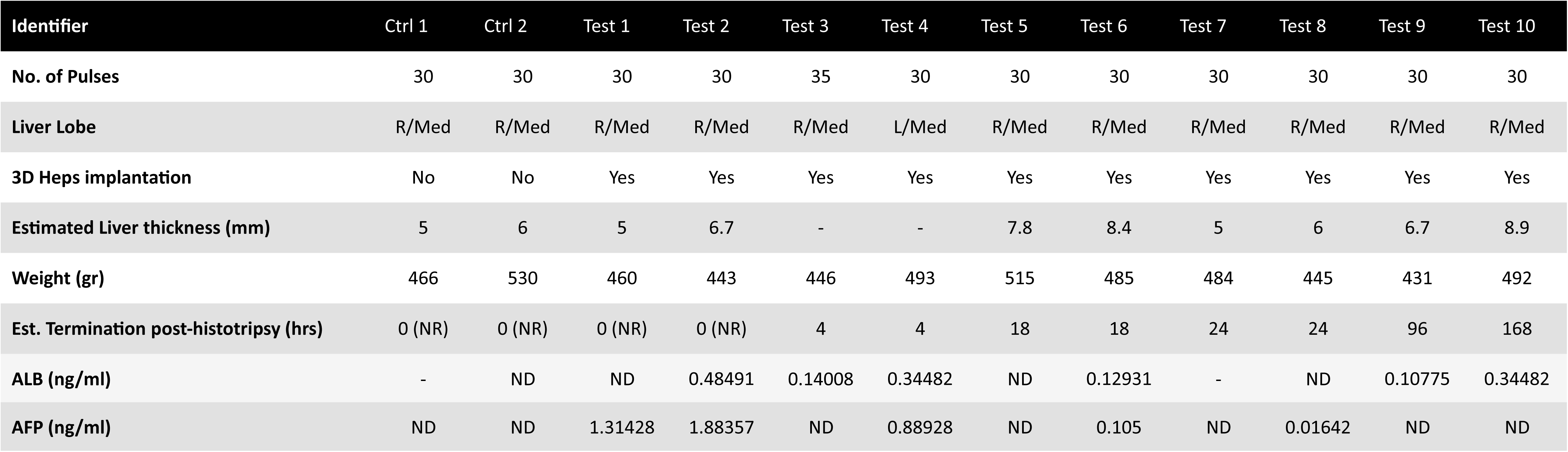
Summary details of feasibility study in Male Sprague–Dawley rats (n=12). **Abbreviations: R/Med, Right Median; L/Med, Left Median; Est, Estimated; NR, Non-recovery; NI, Not Identified; ND, Not-detected.**

