## Supplementary Figure 1 for "The safety and efficacy of Ultrasound Histotripsy and human pluripotent stem cell-derived hepatic spheroid implantation as a potential therapy for treatment of congenital metabolic liver disease: assessment in an immunocompetent rodent model"

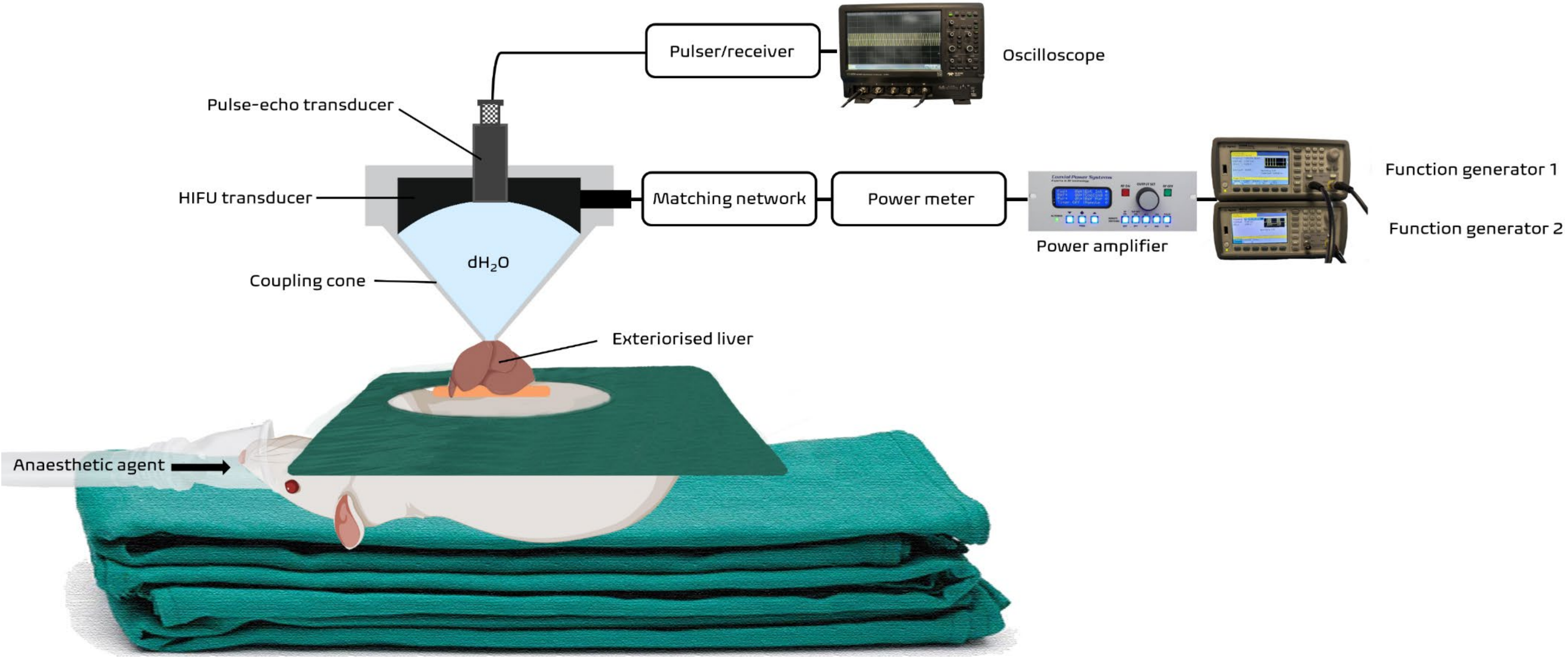


**Supplementary Figure 1: Schematic of histotripsy-mediated cell transplantation. Due to anatomical location and size of the liver in the rat, it was exteriorised following laparotomy before performing histotripsy to create decellularized cavity.**
