## Supplementary Table 1 for "The safety and efficacy of Ultrasound Histotripsy and human pluripotent stem cell-derived hepatic spheroid implantation as a potential therapy for treatment of congenital metabolic liver disease: assessment in an immunocompetent rodent model"

**Supplementary Table 1: Summary details of feasibility study in Male Sprague–Dawley rats (n=12).**

**Abbreviations: R/Med, Right Median; L/Med, Left Median; Est, Estimated; NR, Non-recovery; NI, Not Identified; ND, Not-detected.**

| Identifier | Ctrl 1 | Ctrl 2 | Test 1 | Test 2 | Test 3 | Test 4 | Test 5 | Test 6 | Test 7 | Test 8 | Test 9 | Test 10 |
|---|---|---|---|---|---|---|---|---|---|---|---|---|
| **No. of Pulses** | 30 | 30 | 30 | 30 | 35 | 30 | 30 | 30 | 30 | 30 | 30 | 30 |
| **Liver lobe** | R/Med | R/Med | R/Med | R/Med | R/Med | L/Med | R/Med | R/Med | R/Med | R/Med | R/Med | R/Med |
| **3D Heps implantation** | No | No | Yes | Yes | Yes | Yes | Yes | Yes | Yes | Yes | Yes | Yes |
| **Estimated Liver thickness (mm)** | 5 | 6 | 5 | 6.7 | - | - | 7.8 | 8.4 | 5 | 6 | 6.7 | 8.9 |
| **Weight (gr)** | 466 | 530 | 460 | 443 | 446 | 493 | 515 | 485 | 484 | 445 | 431 | 492 |
| **Est. Termination post-histotripsy (hrs)** | 0 (NR) | 0 (NR) | 0 (NR) | 0 (NR) | 4 | 4 | 18 | 18 | 24 | 24 | 96 | 168 |
| **Cavity distance to liver capsule (mm)** | 0.29 | NI | 0.39 | - | NI | 0.51 | NI | NI | NI | 0.39 | NI | 0.19 |
| **Cavity length (mm)** | 3.39 | NI | 2.89 | 2.01 | NI | 3.28 | NI | NI | NI | 3.54 | NI | 0.5 |
| **Cavity width (mm)** | 1.32 | NI | 0.74 | 0.38 | NI | 0.68-0.77 | NI | NI | NI | 0.83-0.89 | NI | 0.372 |
| **ALB (ng/ml)** | - | ND | ND | 0.48491 | 0.14008 | 0.34482 | ND | 0.12931 | - | ND | 0.10775 | 0.34482 |
| **AFP (ng/ml)** | ND | ND | 1.31428 | 1.88357 | ND | 0.88928 | ND | 0.105 | ND | 0.01642 | ND | ND |
